# Immune Modulation of Allergic Asthma by Early Pharmacological Inhibition of RIP2

**DOI:** 10.1101/2020.06.29.178665

**Authors:** Madelyn H. Miller, Michael G. Shehat, Justine T. Tigno-Aranjuez

## Abstract

Exposure to house dust mite (HDM) is highly associated with the development of allergic asthma. The adaptive immune response to HDM is largely T helper cell type 2 (Th2) dominant and a number of innate immune receptors have been identified which recognize HDM to initiate a Th2 response. Nucleotide-binding Oligomerization Domain-containing Protein 2 (NOD2) is a cytosolic sensor of peptidoglycan which is important for Th2 polarization. NOD2 mediates its signaling through its downstream effector kinase, Receptor-interacting Serine/Threonine Protein Kinase 2 (RIP2). We have previously shown that RIP2 is important in promoting HDM-associated allergic airway inflammation and Th2 immunity. In particular, we demonstrated that the effects of RIP2 were important early in the HDM response, likely within airway epithelial cells. However, the consequences of inhibiting RIP2 during this critical period has not yet been examined. In this study, we pharmacologically inhibited RIP2 activity during the initial exposure to allergen in an acute HDM model of asthma and determined the effect on the subsequent development of allergic airway disease. We show that early inhibition of RIP2 was sufficient to reduce lung histopathology and local airway inflammation while skewing the immune response from Th2 towards Th1. Using a chronic HDM asthma model, we demonstrate that inhibition of RIP2, despite attenuating airway inflammation and airway remodeling, was insufficient to reduce airway hyperresponsiveness. These data demonstrate the potential of pharmacological targeting of this kinase in asthma and support further development and optimization of RIP2 targeted therapies.

## Introduction

Asthma is a chronic inflammatory disease of the airways which affects up to 300 million people worldwide (1). It is characterized by narrowing of the airways, mucus overproduction, airway hyperresponsiveness (AHR) and pathological airway remodeling. Two endotypes (subgroups with similar pathophysiology) are generally recognized for asthma: Type 2 (or type 2 high) and non-type 2 (or type 2 low) (2). Around half of all asthmatics have type 2 inflammation and, of these, the majority suffer from allergic asthma (3). Exposure and sensitization to indoor allergens such as pet dander, cockroaches and dust mites are consistently associated with the development of allergic asthma (4–6). Of these aeroallergens, house dust mites (HDM) are the most frequent sensitizer, with up to 70% of asthmatics showing reactivity to this allergen (7–11).

Type 2 or type-2 helper T cell (Th2) immunity is distinguished by elevated production of the canonical type 2 cytokines IL-4, IL-5 and IL-13 (12). In the pathologic setting of allergic asthma, these cytokines are critical mediators which collectively result in the observed asthmatic symptoms. The large number of therapeutics developed to target these type 2 cytokines and their receptors, or type 2-induced IgE and IgE receptors, as well as clinical translation of such biologics, reflect their importance in the progression of type 2 disease (13). However, the limited indication for such biologics (type 2 asthmatics who have severe disease or high eosinophil counts) as well as the high cost and need for chronic administration, point to the need for novel therapeutic targets which lie upstream of type 2 cytokine production and which have the potential to modulate the type 2 response.

Initiation of type 2 immunity in response to HDM is a process orchestrated by multiple cell types, including epithelial cells, dendritic cells (DCs), group 2 innate lymphoid cells (ILC2s), mast cells, and basophils (12). Likewise, numerous receptors and pathways in innate and structural cells have been implicated in the initial recognition and response to HDM (14). Using knock-out models, our own laboratory has previously demonstrated that Receptor-interacting Serine/Threonine Protein Kinase 2 (RIPK2 or RIP2) is involved in promoting allergic airway inflammation in response to HDM (15). Importantly, multiple lines of evidence, including HDM-induced activation of RIP2 within airway epithelial cells, suggested that the actions of RIP2 were crucial at a very early timepoint. RIP2 is a kinase which mediates signaling downstream of Nucleotide-binding and Oligomerization Domain-Protein 2 (NOD2), a cytosolic receptor for bacterial peptidoglycan (16, 17). Apart from allergic airway inflammation, RIP2 has also been associated with the pathogenesis of various inflammatory diseases including inflammatory bowel disease (IBD), arthritis, and experimental autoimmune encephalomyelitis (EAE) (18–20). These findings have spurred the development and preclinical testing of numerous RIP2 inhbitors as well as initiation of a Phase I clinical trial (21). In the current work, we examine the efficacy of pharmacological inhibition of RIP2 in the setting of a HDM-induced allergic asthma model, and in particular, evaluate the consequences of early and acute inhibition of RIP2. The resulting findings may lend further support for RIP2 as being a viable druggable target in the setting of allergic asthma.

## Materials and Methods

### Mice

C57BL/6J mice (000664) were obtained from Jackson Laboratories (Bar Harbor, Maine) and were housed in specific pathogen free (SPF) and AALAC-accredited animal facility at the UCF Health Science Campus at Lake Nona. Eight week old male and female mice were used for the experiments. All animal protocols were approved by the Univ. of Central Florida IACUC.

### House Dust Mite Asthma Models and GSK Intervention

Lyophilized *Dermatophagoides pteronyssinus* (house dust mite, HDM) was obtained from Greer Laboratories (XPB82D3A2.5, Lot no. 346230) (Lenoir, NC), resuspended in sterile PBS at necessary concentrations, and frozen in aliquots at −20C until use. For all models, irradiated GSK583 chow (0.25g GSK583/ kg chow, C19062701i, Research Diets, New Brunswick, NJ) or irradiated vehicle chow (C13513i, Research Diets, New Brunswick, NJ) was provided ad libitum on days −2 to day 2. This amount will deliver approximately 30 mg/kg/day based on a 3g/day consumption per mouse. On day 3, all mice were switched to vehicle chow.

The acute house dust mite model was conducted as previously published (15). Mice were euthanized on day 1 or day 14 for collection of tissues, depending on the experiment. For the chronic house dust mite model, 40 μL of a 1.5 mg/mL HDM was delivered intratracheally (i.t.) on day 0. Twenty-five microliters of a 0.5 mg/mL HDM mixture was delivered i.t. on days 7-11, and on days 14, 16, 18, 21, 23, and 25. Mice were euthanized on day 28 for collection of tissues. All i.t. installations were performed using isoflurane anesthesia using an otoscope, endotracheal tube, guide wire, and gas-tight syringe for delivery of HDM.

### Histopathological scoring

Lung tissue was harvested from euthanized mice for histology. Tissue was fixed in 10% buffered formalin and sent to AML labs (St. Augustine, FL) for paraffin embedding, sectioning, and Hematoxylin & Eosin (H&E), Periodic Acid-Schiff (PAS), and Masson’s Trichrome staining to assess total inflammation, mucus production and collagen deposition/fibrosis, respectively. All sections were scored blindly using a modified histopathological scoring system as previously published (15, 22). The maximum combined score for lung inflammation was 16, the maximum combined score for mucus production was four, and the maximum combined score for trichrome staining was 6 (0 being least severe and the maximum score being most severe). For the chronic model, an alternative scoring system to assess chronic airway remodeling was used, for a maximum combined score of 22. The following parameters were assessed for incidence throughout the section: epithelial folding/distortion (0-2), goblet cell metaplasia (0-2), smooth muscle cell hypertrophy (0-2), subepithelial fibrosis (0-2), peribronchial trichrome blue staining (0-3), and parenchymal collagen deposition (0-3). In addition, the following parameters were assessed for severity: epithelial folding/distortion (0-2), goblet cell metaplasia (0-2), smooth muscle cell hypertrophy (0-2), and subepithelial fibrosis (0-2).

### Flow cytometry

On day 14, bronchoalveolar lavage (BAL) and a portion of the right lung lobe were collected and processed as previously described. To assess cellular infiltrate, one million cells of lung or BAL were stained using a previously described antibody cocktail and gating strategy. Lung tissue was collected, processed, stimulated with PMA (5ng/mL) and ionomycin (500ng/mL) and stained for cell surface antigens and intracellular cytokines. The stimulation, fixing and permeabilization conditions, as well as the antibody cocktail and gating strategy used were similar to that described previously (15). All samples were acquired using a Novocyte flow cytometer (ACEA Biosciences, San Diego, CA) and analyzed using the NovoExpress Software.

### Lung homogenate cytokine analysis

A portion of the right lobe of the lung was homogenized in T-PER buffer with protease inhibitors (ThermoFisher, Waltham, MA). Samples were normalized to 0.5 mg/mL protein in Legendplex assay buffer (Biolegend, San Diego, CA) using a Bradford Assay (BioRad, Hercules, CA). A 13-plex bead-based immunoassay was performed using the manufacturer’s instructions (Legendplex Th Panel, Biolegend, San Diego, CA). All samples were acquired using Novocyte flow cytometer (ACEA Biosciences, San Diego, CA) and analyzed using the Legendplex software (v8, Vigenetech, Carlisle, MA).

### Serum antibody analysis

Mice were subjected to cardiac puncture for collection of blood, which was clotted for 30 minutes in serum separator tubes prior to centrifugation for 10 mins at 4C for collection of serum. Serum was diluted 1:10 in assay diluent. Murine anti-HDM IgG1 and IgG ELISA kits were obtained from Chrondrex, Inc. (Redmont, WA) and used as directed by the manufacturer. Assays were developed and analyzed as previously described (15).

### Flexivent analysis for airway hyperresponsiveness

On day 28, mice were anesthetized using a cocktail of Ketamine/Xylazine/Acepromazine (65 mg/kg, 13 mg/kg, 2mg/kg, respectively). Once appropriate plane of anesthesia was achieved, mice were cannulated intratracheally and connected to a FlexiVent mechanical ventilator (Scireq). Basal lung mechanics and airway hyperresponsiveness to increasing doses of inhaled aerosolized methacholine (0, 3.125, 6.25, 12.5, 25, 50, and 100mg/mL) was measured using pre-set scripts. Deep Inflation, SnapShot-150, Quick Prime-3, and PVs-P perturbations were collected at baseline three times. For each dose, SnapShot-150 and Quick Prime-3 perturbations were collected 12 times. Analysis was conducted using the FlexiWare 7 Software (Scireq).

### Statistical analysis

Statistical analysis was conducted using GraphPad Prism. Significance levels were fixed at 5% for each measured response. Figure legends indicate specific tests used for analysis of each dataset, number of animals per group, and number of times the experiment was repeated. Bar heights indicate means and error bars indicate SEM.

## Results

### Early inhibition of RIP2 reduces lung pathology in mice subjected to an acute house dust mite asthma model

We have previously shown that genetic loss of RIP2 attenuates lung pathology in response to HDM exposure (15). In the prior study, we had additional evidence to suggest that the effects of RIP2 were mediated at a very early timepoint using complete genetic knockouts where RIP2 was absent throughout the model. In the absence of conditional, inducible, or floxed RIP2 strains, in the current study, we chose to investigate the role of RIP2 specifically during the early reponse to HDM through pharmacological inhibition. We subjected mice to an acute HDM model of asthma and provided a selective RIP2 inhibitor (GSK583) via chow at a dose of 30 mg/kg/day on the 5 days during and surrounding the initial exposure to HDM (days −2 to 2, Figure 1A). The specificity, mechanism and in vivo inhibitory activity of GSK583 have been previously reported elsewhere (23). On day 14 of this model, mice were euthanized and lungs were harvested and fixed for histological examination. Paraffin-embedded sections were stained using Hematoxylin & Eosin (H&E), Periodic Acid Schiff (PAS), or Trichrome stains, then scored blindly. We used a combination of parameters to create an inflammatory index (maximum score of 16) comprised of individual scores for bronchoarterial inflammation, amuscular blood vessel inflammation, inter-alveolar space inflammation, pleural inflammation, and pulmonary vein inflammation. Early inhibition of RIP2 in GSK583-treated mice was sufficient to reduce overall inflammation compared to their vehicle-treated counterparts (Figure 1B, with corresponding graph in Figure 1C). Similarly, this early inhibition of RIP2 using GSK583 was sufficient to reduce mucus production as seen with PAS staining (magenta in PAS stain in Figure 1B, with corresponding graph in Figure 1D) and collagen deposition as seen with trichrome staining (blue in trichrome stain in Figure 1B, with corresponding graph in Figure 1G) compared to vehicle-treated mice. In addition, GSK583-treated mice demonstrated reduced lumen narrowing (increased lumen area) (Figure 1E), and decreased epithelial thickness (Figure 1F) compared to vehicle-treated mice subjected to an acute HDM asthma model. Collectively, these data show that RIP2 inhibition using GSK583 during the initial exposure to HDM is sufficient to improve lung pathology in an acute house dust mite asthma model.

**Figure 1.**
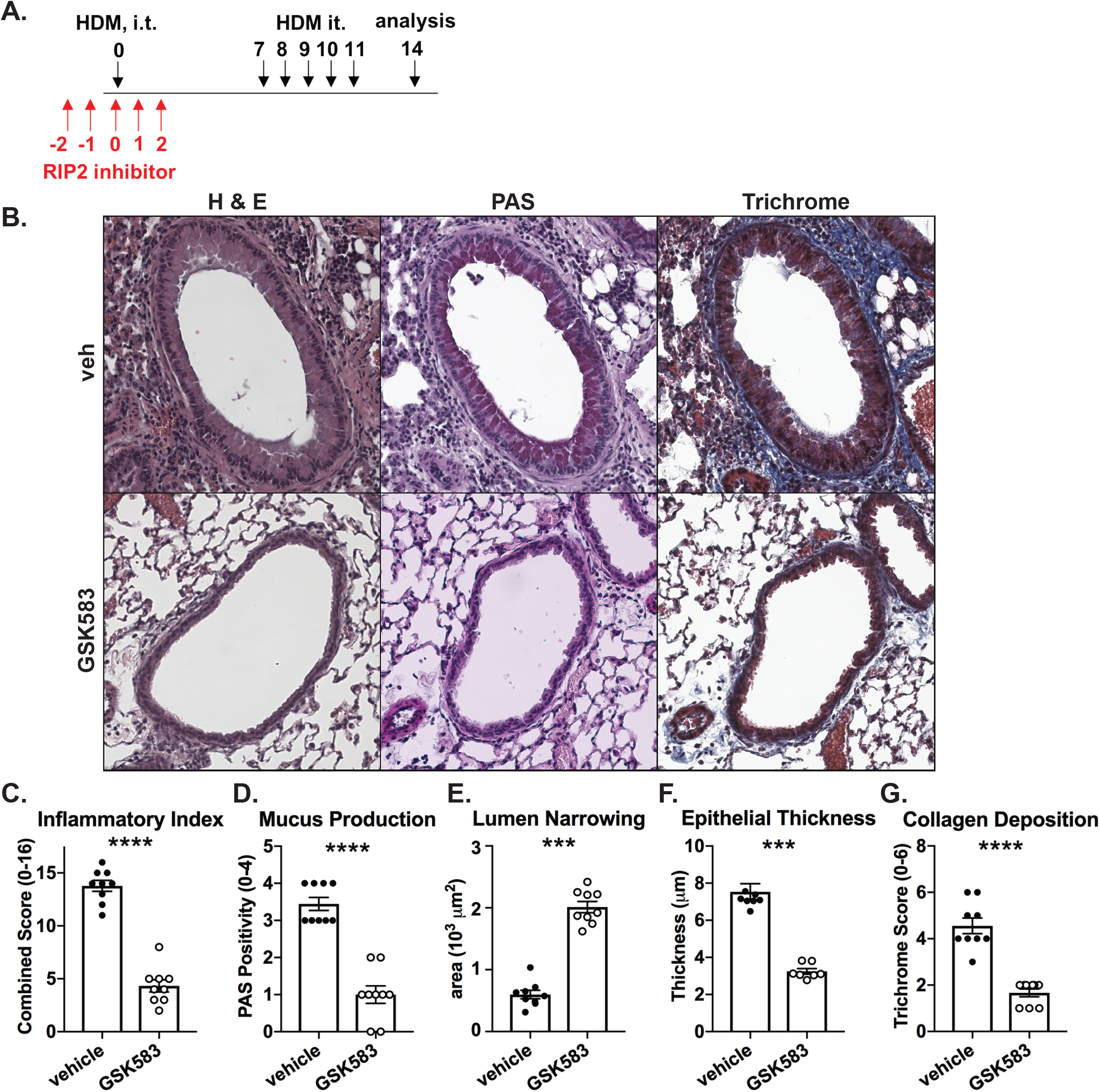
Early prophylactic inhibition of RIP2 reduces lung pathology in an acute house dust mite (HDM) model of asthma. A.) WT C57BL/6 mice were subjected to an acute HDM asthma model and were administered either regular chow or chow containing RIP2 inhibitor (GSK583 at 30mg/kg/day) for the 5 days during and surrounding the initial exposure to HDM (red arrows). Mice were euthanized on day 14 and lungs were harvested, fixed in 10% formalin, and embedded in paraffin. B.) Consecutive sections were stained with either H&E, PAS, or Trichrome. Severity of lung pathology was scored based on C.) inflammatory index, D.) mucus production, E.) lumen narrowing, F.) epithelial thickness, and G.) collagen deposition and was performed by a blinded observer. Data are presented as scatterplots with bars where bar heights represent means ± SEM. Filled in circles represent individual datapoints for vehicle/control chow-treated mice and open circles represent individual datapoints for GSK583-treated mice. Data are pooled from 3 independent experiments for a total of n=9 mice per group. Statistical analysis was performed using an unpaired, two-tailed Student’s t-test. *** = *p*<0.001, **** = *p*<0.0001.

### Early prophylactic inhibition of RIP2 reduces eosinophilia in a HDM model of asthma

To assess the effect of early pharmacological inhibition of RIP2 in an HDM asthma model on the recruitment of inflammatory cells to the airway, we harvested bronchoalveolar lavage (BAL) and lung cells and stained these using a panel of antibodies which would allow discrimination of hematopoietic cells, lymphocytes, neutrophils, alveolar macrophages, and eosinophils when subjected to flow cytometric analysis. Figure 2A shows that mice undergoing pharmacological inhibition of RIP2 during initial exposure to HDM allergen (GSK583-treated mice) have significantly reduced numbers of CD45^+^ cells and eosinophils recovered from the BAL (corresponding representative gating strategies shown in Figure 2B). There was a trend for decreased numbers of lymphocytes and neutrophils recovered in the GSK583-treated BAL compared to vehicle-treated mice; although this difference was not significant. Similar to the BAL, early inhibition of RIP2 led to significantly reduced percentages of CD45^+^ cells and eosinophils recruited to the lung compared to vehicle treated mice (Figure 2C, with corresponding representative gating strategies shown in Figure 2D). Overall, these results indicate that early inhibition of RIP2 can reduce eosinophilia during an acute house dust mite asthma model.

**Figure 2.**
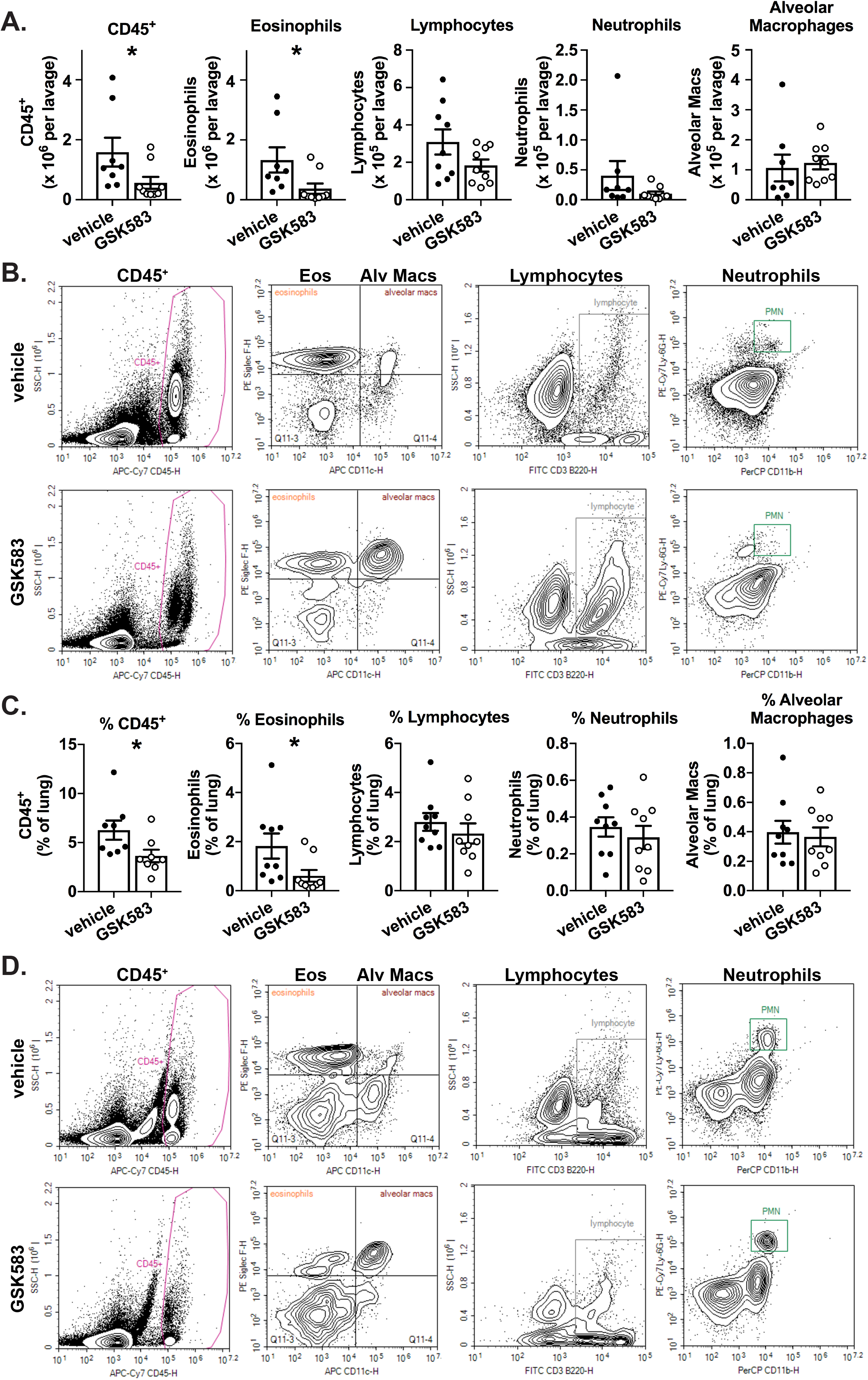
Early prophylactic inhibition of RIP2 reduces eosinophilia in an acute HDM model of asthma.. WT C57BL/6 mice were subjected to an acute HDM asthma model and were administered either regular chow or chow containing RIP2 inhibitor (GSK583) as indicated in Fig 1A. On day 14, mice were euthanized and bronchoalveolar lavage (BAL) cells or dissociated lung cells were isolated, stained for immune cell markers, and subjected to flow cytometric analysis. A). The total number of each cellular population within the BAL for vehicle compared to GSK583-treated mice. B.) Flow cytometry gating strategy for obtaining BAL cell counts in A.). C.) The percentage of each cellular population in the lung is shown for vehicle compared to GSK583-treated mice. D.) Flow cytometry gating strategy for obtaining lung cell percentages in C.). Data are presented as scatterplots with bars where bar heights represent means ± SEM. Filled in circles represent individual datapoints for vehicle/control chow-treated mice and open circles represent individual datapoints for GSK583-treated mice. Data are pooled from 3 independent experiments for a total of n=9 mice per group. Statistical analysis was performed using an unpaired, two-tailed Student’s t-test. *= *p*<0.05.

### Early pharmacological inhibition of RIP2 downregulates Th2 immunity and upregulates Th1 immunity in an acute HDM model of asthma

The adaptive immune response during an HDM asthma model is largely Th2 and Th17 dominated. To assess the effects of early pharmacological inhibition of RIP2 on the resulting adaptive immune response, we collected lung cells and mediastinal lymph nodes at day 14 of the acute HDM asthma model, stimulated these with PMA+ionomycin and performed intracellular cytokine staining to assess production of Th1, Th2, or Th17-associated cytokines. Interestingly, while we observed a reduction in Th2 (CD4^+^IL-4^+^IL-5^+^) cells within the lung in GSK583-treated mice as expected from the decrease in eosinophilia, there was also a significant increase in the percentage of Th1 (CD4^+^IFN-γ^+^) cells in the lung without a remarkable change in Th17 (CD4^+^IL-17^+^) cell numbers (Figure 3A). Corresponding representative gating strategies are shown in Figure 3B. A similar trend was found when ICS was performed on lung draining lymph nodes (Supplemental Fig 1.).

**Figure 3.**
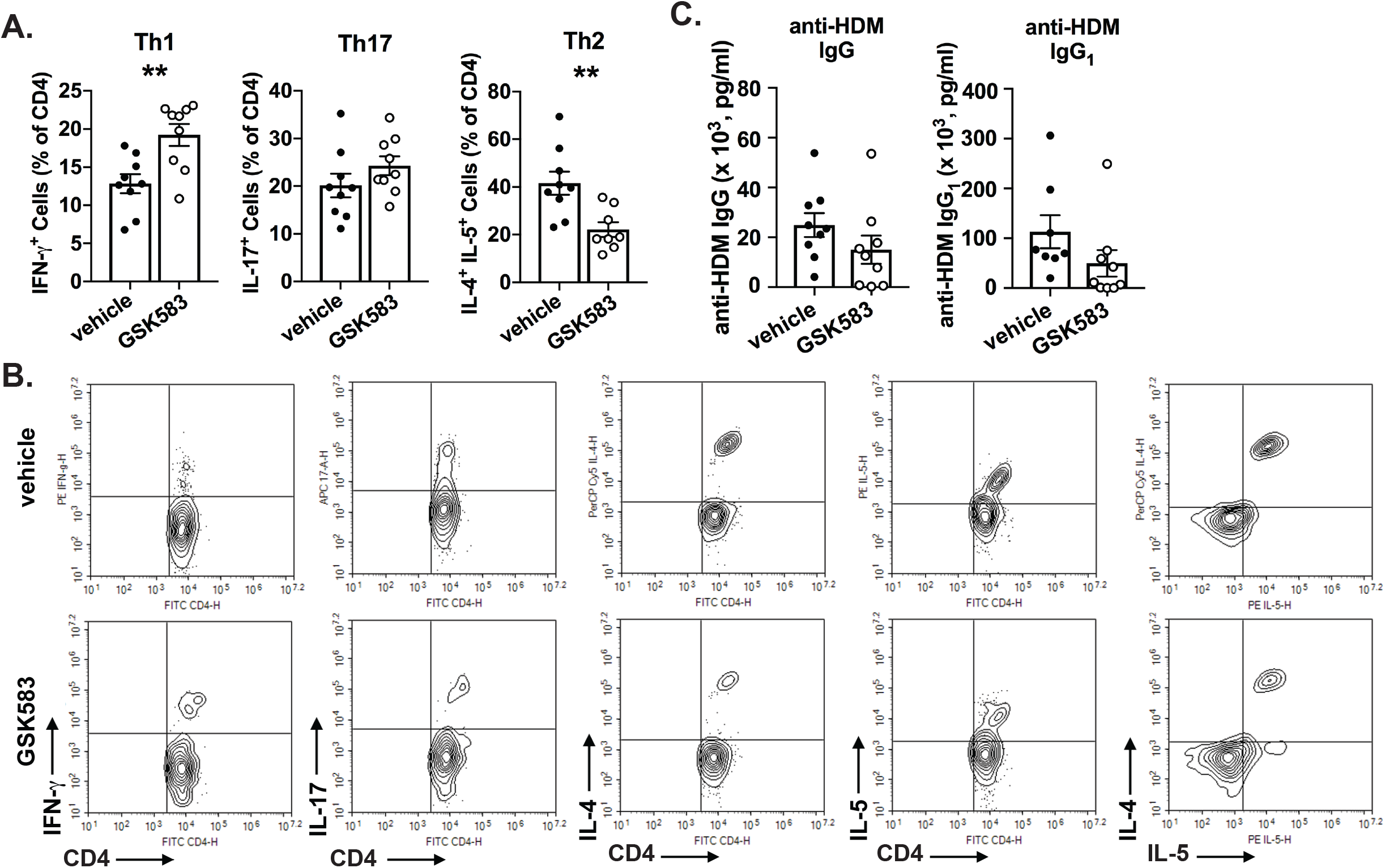
Pharmacological inhibition of RIP2 downregulates Th2 immunity and upregulates Th1 immunity in a HDM model of asthma. WT C57BL/6 mice were subjected to an acute HDM asthma model and were administered either regular chow or chow containing RIP2 inhibitor (GSK583) as indicated in Fig 1A. On day 14, mice were euthanized and dissociated lung cells were stimulated with PMA and ionomycin and subjected to intracellular cytokine staining (ICS). A.) The percentage of CD4^+^IFN-γ^+^(Th1), CD4^+^IL-17^+^(Th17), or CD4^+^IL-4^+^IL-5^+^(Th2) are shown for vehicle compared to GSK583-treated mice. B.) Flow cytometry gating strategy for ICS percentages shown in A.). C.) Levels of HDM-specific total IgG and IgG1 antibodies in the serum of vehicle chow or RIP2 inhibitor (GSK583) chow-treated mice subjected to an acute HDM asthma model as measured by ELISA. Data are presented as scatterplots with bars where bar heights represent means ± SEM. Filled in circles represent individual datapoints for vehicle/control chow-treated mice and open circles represent individual datapoints for GSK583-treated mice. Data are pooled from 3 independent experiments for a total of n=9 mice per group. Statistical analysis was performed using an unpaired, two-tailed Student’s t-test. **= *p*<0.01.

Furthermore, we also assessed the effect of early and acute RIP2 inhibition on the production of HDM-specific antibodies. We have previously shown that on the C57BL/6J background (and without the use of an additional adjuvant such as alum), the HDM asthma model leads primarily to an increase in the production of HDM-specific total IgG and the Th2-associated IgG1 subclass of antibodies (15). Changes in levels of these antibodies in the serum of GSK583 or vehicle-treated animals suggested a trend towards decreased levels of HDM-specific total IgG and IgG1; however, this was not statistically significant (Figure 3C).

To additionally examine the local adaptive immune response within the lung, we performed a bead-based multiplex assay with lung homogenates from GSK583 and vehicle-treated mice (Figure 4). This assay measures cytokines collectively secreted by the major Th lineages (Th1, Th2, Th9, Th17, Th22 and T follicular cells). These data indicate that there was a significant reduction in the levels of IL-4 and IL-5 in the lungs of GSK583-treated mice compared to vehicle-treated mice on an HDM asthma model without appreciable alteration of any of the other 11 Th cytokines tested (Figure 4). Collectively, these data indicate that pharmacological inhibition of RIP2 during the initial exposure to HDM is enough to reduce local Th2-cytokines and to skew the adaptive immune response from Th2 towards Th1 immunity.

**Figure 4.**
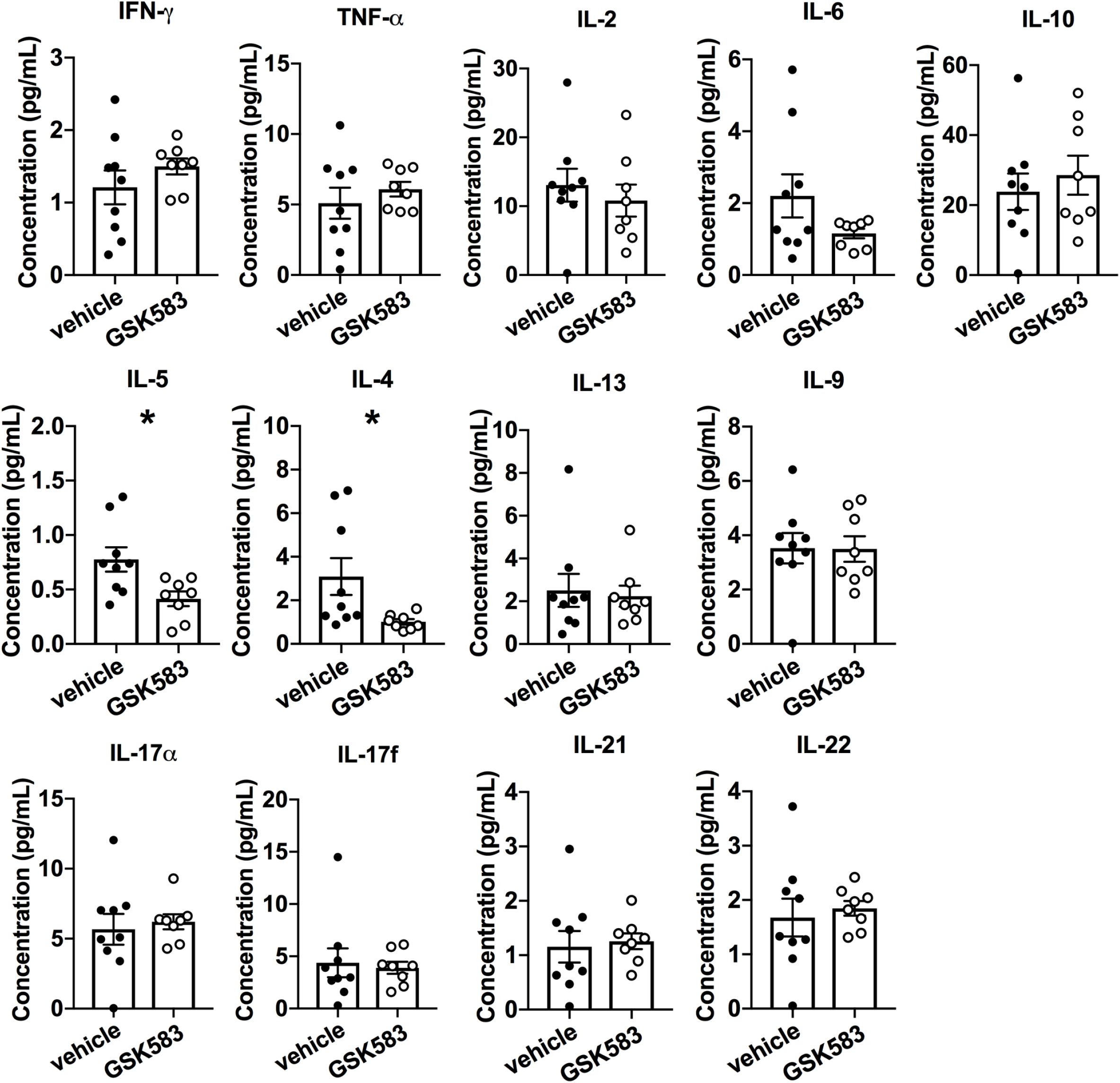
Early in vivo inhibition of RIP2 reduces Th2 associated cytokines in the lung. WT C57BL/6 mice were subjected to an acute HDM asthma model and were administered either regular chow or chow containing RIP2 inhibitor (GSK583) as indicated in Fig 1A. On day 14, mice were euthanized and lung tissue was collected and lysed in protein extraction buffer containing inhibitors. Supernatants were subjected to multiplexed bead-based immunoassay for measurement of various Th-associated cytokines. Data are presented as scatterplots with bars where bar heights represent means ± SEM. Filled in circles represent individual datapoints for vehicle/control chow-treated mice and open circles represent individual datapoints for GSK583-treated mice. Data are pooled from 3 independent experiments for a total of n=9 mice per group. Statistical analysis was performed using an unpaired, two-tailed Student’s t-test. Statistical analysis was performed using an unpaired, two-tailed Student’s t-test. *= *p*<0.05.

### Early inhibition of RIP2 in mice undergoing a chronic HDM asthma model exhibits disparate effects on lung pathology and airway hyperresponsiveness (AHR)

Given that acute models of asthma do not recapitulate many important features of this chronic disease such as airway remodeling and airway hyperresponsiveness (AHR), we also wanted to utilize a chronic HDM asthma model to determine whether early pharmacological inhibition of RIP2 had beneficial effects which would still be evident even at the later stages of this disease. As such, we treated mice with GSK583 or vehicle chow for 5 days around the time of initial exposure to HDM allergen and continued to expose them to HDM for a period of 4 weeks as depicted in Figure 5A. On day 28 of the model, mice were euthanized and lungs were harvested, fixed, paraffin-embedded and stained for histological examination. H&E, PAS, and trichrome staining was performed and sections were scored by a blinded observer. Similar to the acute model, early inhibition of RIP2 using GSK583 was sufficient to reduce the total inflammatory index (Figure 5B, with corresponding graph in Figure 5C) and mucus production as shown by PAS positivity (magenta in PAS staining in Figure 5B, with corresponding graph in Figure 5E) compared to vehicle treated mice when mice were subjected to a chronic HDM model. In addition to evaluating inflammation, we also assessed airway remodeling. Using a combination of parameters encompassing severity and incidence of epithelial folding/distortion, goblet cell metaplasia, smooth muscle cell hypertrophy, subepithelial fibrosis, peribronchial trichrome blue staining, and parenchymal trichrome blue staining, we report an airway remodeling index (maximum score of 22). Early inhibition of RIP2 using GSK583 exhibited significantly reduced airway remodeling compared to vehicle-treated mice (11.22 ± 0.88 compared to 21 ± 0.29) when mice were subjected to a chronic model of asthma (Figure 5B, with corresponding graph in Figure 5D). These results indicate that by several histological parameters, early inhibition of RIP2 is enough to improve lung pathology and airway remodeling even during a chronic model of asthmatic disease.

**Figure 5.**
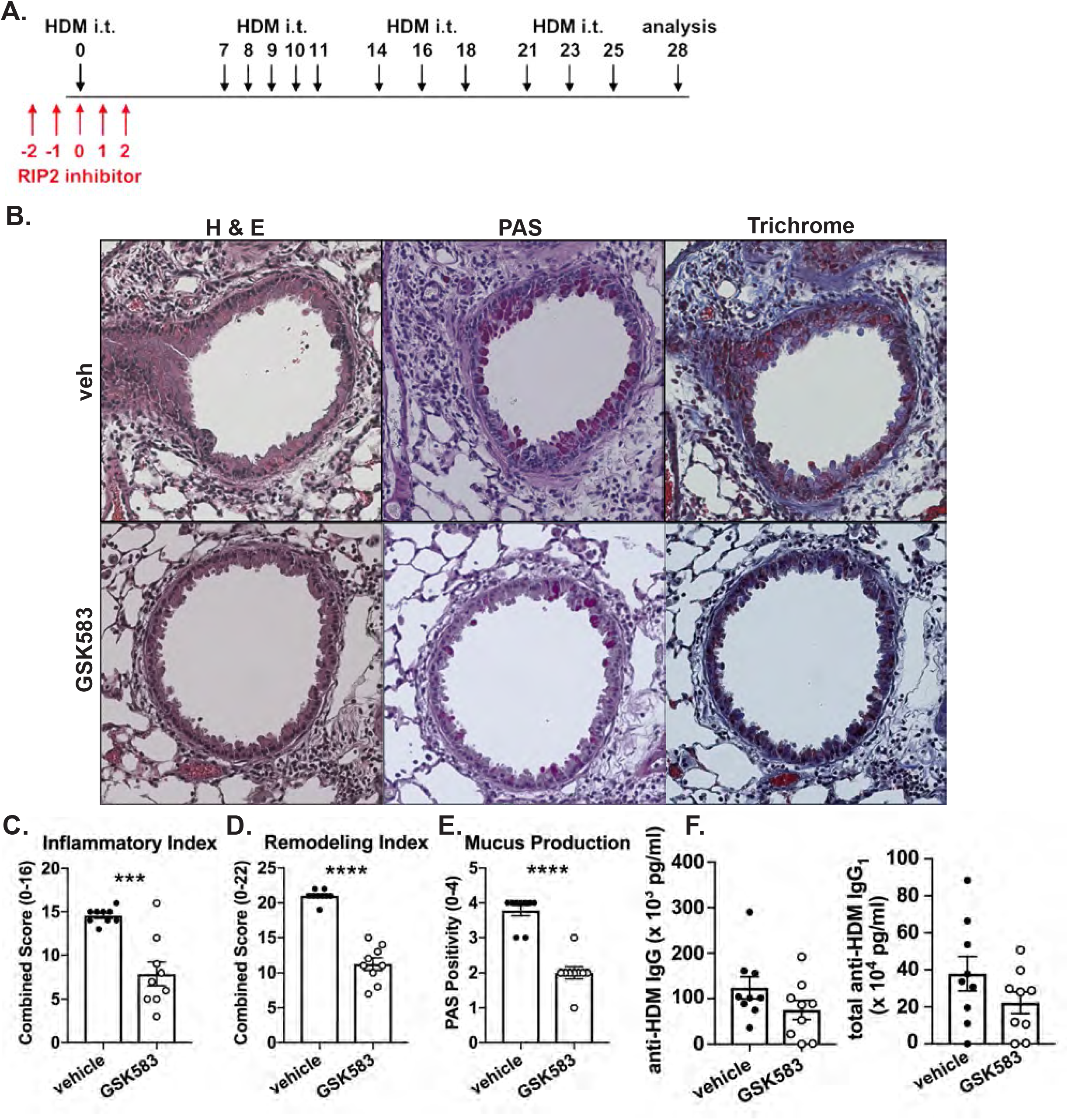
Mice undergoing a chronic HDM asthma model exhibit improved lung pathology with early RIP2 inhibition but minimal effects on HDM-specific antibody responses. A.) WT C57BL/6 mice were subjected to a chronic HDM asthma model and were administered either regular chow or chow containing RIP2 inhibitor (GSK583 at 30mg/kg/day) for the 5 days during and surrounding the initial exposure to HDM (red arrows). On day 28, mice were euthanized and lungs were harvested, fixed in 10% formalin, and embedded in paraffin. B.) Consecutive sections were stained with either H&E, PAS, or Trichrome. Severity of lung pathology was blindly scored based on C.) inflammatory index, D.) remodeling index and E.) mucus production. F.) Levels of HDM-specific total IgG and IgG1 antibodies in the serum of vehicle chow or RIP2 inhibitor (GSK583) chow-treated mice subjected to a chronic HDM asthma model as measured by ELISA. Data are presented as scatterplots with bars where bar heights represent means ± SEM. Filled in circles represent individual datapoints for vehicle/control chow-treated mice and open circles represent individual datapoints for GSK583-treated mice. Data are pooled from 3 independent experiments for a total of n=9 mice per group. Statistical analysis was performed using an unpaired, two-tailed Student’s t-test. ***= *p*<0.001, ****= *p*<0.0001

In addition, we assessed whether treatment with GSK583 would lead to deficient HDM-specific antibody responses in the chronic HDM asthma model. Similar to the acute model, there was a trend for reduced HDM-specific total IgG and IgG1 antibody levels in GSK583-treated mice compared to vehicle-treated mice; however, this was not significant (Figure 5F).

Lastly, to determine whether pharmacological inhibition of RIP2 during initial exposure to allergen affected AHR, mice treated with either GSK583 or vehicle chow underwent a chronic HDM asthma model. On day 28, mice were anesthetized, cannulated, connected to the FlexiVent ventilator and subjected to various mechanical perturbations in a customized asthma challenge script which included administration of nebulized methacholine at increasing doses in between measruments. Resistance of the total respiratory system (Rrs), central airway Newtonian resistance (RN), Elastance of the respiratory system (Ers), tissue damping (Max G), and tissue elastance (Max H) were calculated for each mouse at each dose of methacholine. No significant changes in R_rs_ (Figure 6A) or R_N_ (Figure 6B) were observed with GSK583 treatment compared to vehicle treatment. Ers (Figure 6C), Max G (Figure 6D), and Max H (Figure 6E) were slightly decreased in GSK583-treated mice compared to vehicle-treated mice at a the highest dose of MCh (100mg/mL); however, these changes were not significant. Overall, these results indicate that early inhibition of RIP2, while sufficient to reduce lung pathology and airway remodeling, is by itself not sufficient to decrease AHR in a chronic house dust mite asthma model.

**Figure 6.**
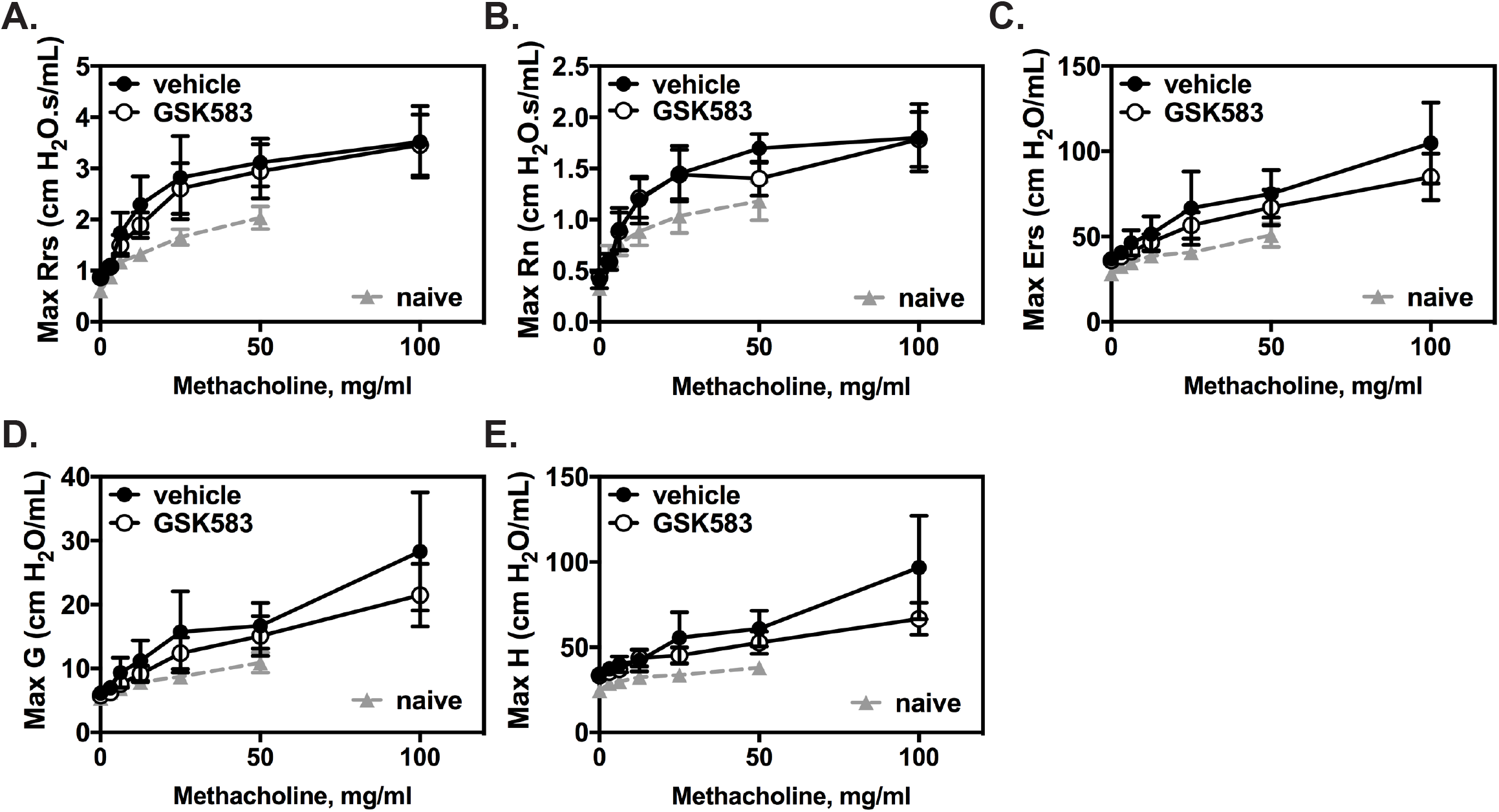
GSK583-treated mice did not exhibit changes in HDM-induced airway hyperresponsiveness. WT C57BL/6 mice were subjected to a chronic HDM asthma model and were administered either regular chow or chow containing RIP2 inhibitor (GSK583) as indicated in Fig 5A. On day 28, mice were anesthetized, intratracheally cannulated, and connected to a FlexiVent mechanical ventilator. Baseline lung function was obtained, after which, mice were subjected to increasing doses of methacholine to measure airway hyperresponsiveness. Dose dependent changes in A.) Resistance of the respiratory system (Rrs), B.) Resistance of the central airways (Rn), C.) Elastance of the respiratory system (Ers), D.) Tissue damping and E.) Tissue elastance (H) are shown. Lines connected by filled in circles represent vehicle/control chow-treated mice and lines connected by open circles represent individual datapoints for GSK583-treated mice. Data are pooled from 2 independent experiments for a total of n=5-6 mice per group. The area under the curve (AUC) was calculated for each treatment and an unpaired, two-tailed Student’s t-test was performed comparing the two.

## Discussion

Currently, the only disease modifying treatment available for asthmatics is allergen immunotherapy (AIT) (24). AIT has a long history of efficacy primarily in the form of subcutaneous immunotherapy (SCIT or “allergy shots”) and, more recently, as sublingual formulations (SLIT). The proposed mechanisms for the effectiveness of AIT include induction of T cell anergy, induction of regulatory T cells (Tregs) or immune deviation (25). The result is the production of antibodies of the IgG4 or IgG isotype with the ability to block allergen specific IgE. Although this treatment has been available for some time, it is generally underused due to lack of standardization (allergen preparation, potency, delivery system, route of delivery) and is contraindicated for patients with uncontrolled asthma (25).

Recently, the FDA approved a Phase 3 clinical trial for use of the anti-TSLP monoclonal antibody, tezepelumab in adults with severe uncontrolled asthma (26). TSLP is an alarmin produced by the airway epithelium in response to allergen stimulation (27). Co-administration of TSLP with HDM allergens during T cell priming has been shown to promote Th2 responses, indicating a role for TSLP in the development of Th2 immunity (28). Blocking TSLP or its receptor in ovalbumin or HDM models of asthma led to a inversion of the increased Th2/Th1 ratio present in allergic airway inflammation (29). These findings, combined with the fact that TSLP therapies have progressed to Phase 3 clinical trials, demonstrate the potential for therapeutic strategies which target the early response to aeroallergens and which modulate the response away from type 2 immunity.

Our previous study identified RIP2 as a kinase important in promoting not only the Th2 response to HDM, but also the accompanying airway eosinophilia and lung pathology (15). Loss of RIP2 inhibited the early HDM-induced pro-inflammatory chemokine response in the lung and in airway epithelial cells without influencing ILC2 expansion. Although we had previously implicated RIP2 in this process, RIP2 was absent allthroughout the asthma model (RIP2 KOs) and early effects of RIP2 within innate or structural cells could not be distinguished from any potential late effects on adaptive immunity. In this current study, we are able to verify and isolate the importance of RIP2 during the initial period leading to the establishment of T cell immunity by pharmacological inhibition of RIP2 during a crucial window (initial allergen exposure). We demonstrate that early acute inhibition of is beneficial and immune modulating and that the effects on cellular inflammation and lung pathology (airway remodeling) persist long after treatment is discontinued. Although the lack of efficacy of RIP2 inhibition on airway hyperresponsiveness is somewhat disappointing (no change in central airway resistance, RN), the trend in the reduction of tissue damping and elastance are suggestive for a potential role of RIP2 in influencing localized peripheral effects in the lung tissue, changes which may be more apparent if using a different genetic background or sensitization procedure (alum adjuvanted). We performed the experiments as presented to illustrate the temporal importance of RIP2 activity during the early response to HDM which we demonstrate influences the later establishment of Th2 immunity. In practicality, it would be difficult to capture the period at which sensitization to allergens occurs. Thus, in future work, we would additionally like to assess the usefulness of such a treatment using a therapeutic regimen in an established model of allergic asthma. We suspect that although a similar benefit in attenuation of allergic airway inflammation and Th2 immunity may be observed when RIP2 inhibitors are administered therapeutically, there would be additional mechanisms at play. Given the widespread expression and importance of RIP2 not only within epithelial cells but also in various immune cell subsets present during the effector phase, we consider this beyond the scope of the current study.

Use of an immunomodulatory small molecule inhibitor (SMI) needing only intermittent or brief adminstration for treatment of asthma would be very appealing. A short therapeutic regimen would additionally reduce any potential concerns about immunosuppression given the fact that RIP2 does carry out many host protective functions. As a potential therapeutic target, RIP2 has garnered a great deal of interest as beneficial effects of genetic loss or pharmacological inhibition of this kinase has been demonstrated in models of inflammatory bowel disease (19, 30), *ex vivo* sarcoidosis tissues (31), a model of multiple sclerosis (18, 32) and various cancers (33, 34). This work is the first to demonstrate the efficacy and modulatory activity of a RIP2 targeting compound using a clinically relevant model of allergic asthma. Numerous inhibitors have both been discovered and developed for modulation of RIP2 activity (23, 35–43). The RIP2 inhibitor utilized in this study, GSK583, has been reported to be highly selective and exhibits strong potency even *in vivo* (23, 39). Numerous reports have emerged recently regarding the mechanism of action of such inhibitors in attenuating the downstream inflammatory signaling emanating from RIP2. Although many RIP2-targeted therapies strongly inhibit the kinase function of RIP2, the ability of RIP2 to bind XIAP and undergo ubiquitination rather than its kinase activity *per se* appears to be crucial for propagation of NOD2 inflammatory signaling (39, 40). GSK583 has been demonstrated to disrupt XIAP:RIP2 binding and prevent RIP2 ubiquitination and optimized derivatives of this compound (GSK2983559) briefly entered into human clinical trials in 2017 (21). However, this trial was terminated in 2019 due to “non clinical toxicology findings and reduced safety margins” (21). The data presented in this current study, demonstrating a clear benefit of acute and early RIP2 inhibition for attenuation of allergic asthma and modulation of Th2 immunity, will hopefully provide the needed rationale and support for continued development of what will likely be very useful compounds for asthma and other inflammatory diseases.

## Acknowledgments

We would like to thank Pamela Haile, Bart Votta, Allison Beal and John Lich for their assistance in selection of the GSK583 RIP2 inhibitor, determination of the appropriate dose in chow and for providing the compound for use in this study. We additionally would like to thank them for helpful comments during revision of the manuscipt.

We would like to thank UCF Core facilities for use of the GentleMacs tissue dissociator and Madelyn’s thesis committee for discussions regarding her work.

## Author Contributions

M.H.M. performed the experiments, acquired data, and contributed to the writing of the manuscript

M.G.S. performed the experiments, acquired data, and contributed to the writing of the manuscript

J.T.T-A. conception and design of the project, performed the experiments, acquired data, contributed to the writing of the manuscript, approved the final version

## References

1. Organization WH. Global Health Estimates 2016: Deaths by Cause, Age, Sex, by Country and by Region, 2000-2016. Geneva: WHO Press; 2018.

2. Kuruvilla ME, Lee FE, Lee GB. Understanding Asthma Phenotypes, Endotypes, and Mechanisms of Disease. Clin Rev Allergy Immunol. 2019;56(2):219–33.

3. Fahy JV. Type 2 inflammation in asthma--present in most, absent in many. Nat Rev Immunol. 2015;15(1):57–65.

4. Gaffin JM, Phipatanakul W. The role of indoor allergens in the development of asthma. Current opinion in allergy and clinical immunology. 2009;9(2):128–35.

5. Illi S, von Mutius E, Lau S, Niggemann B, Gruber C, Wahn U, et al. Perennial allergen sensitisation early in life and chronic asthma in children: a birth cohort study. Lancet. 2006;368(9537):763–70.

6. Kusel MM, de Klerk NH, Kebadze T, Vohma V, Holt PG, Johnston SL, et al. Early-life respiratory viral infections, atopic sensitization, and risk of subsequent development of persistent asthma. J Allergy Clin Immunol. 2007;119(5):1105–10.

7. Sporik R, Chapman MD, Platts-Mills TA. House dust mite exposure as a cause of asthma. Clinical and experimental allergy : journal of the British Society for Allergy and Clinical Immunology. 1992;22(10):897–906.

8. Sears MR, Herbison GP, Holdaway MD, Hewitt CJ, Flannery EM, Silva PA. The relative risks of sensitivity to grass pollen, house dust mite and cat dander in the development of childhood asthma. Clinical and experimental allergy : journal of the British Society for Allergy and Clinical Immunology. 1989;19(4):419–24.

9. Sporik R, Holgate ST, Platts-Mills TA, Cogswell JJ. Exposure to house-dust mite allergen (Der p I) and the development of asthma in childhood. A prospective study. N Engl J Med. 1990;323(8):502–7.

10. Celedon JC, Milton DK, Ramsey CD, Litonjua AA, Ryan L, Platts-Mills TA, et al. Exposure to dust mite allergen and endotoxin in early life and asthma and atopy in childhood. J Allergy Clin Immunol. 2007;120(1):144–9.

11. Tovey ER, Almqvist C, Li Q, Crisafulli D, Marks GB. Nonlinear relationship of mite allergen exposure to mite sensitization and asthma in a birth cohort. J Allergy Clin Immunol. 2008;122(1):114–8, 8 e1-5.

12. Lambrecht BN, Hammad H, Fahy JV. The Cytokines of Asthma. Immunity. 2019;50(4):975–91.

13. Doroudchi A, Pathria M, Modena BD. Asthma biologics: Comparing trial designs, patient cohorts and study results. Ann Allergy Asthma Immunol. 2020;124(1):44–56.

14. Huang FL, Liao EC, Yu SJ. House dust mite allergy: Its innate immune response and immunotherapy. Immunobiology. 2018;223(3):300–2.

15. Miller MH, Shehat MG, Alcedo KP, Spinel LP, Soulakova J, Tigno-Aranjuez JT. Frontline Science: RIP2 promotes house dust mite-induced allergic airway inflammation. Journal of leukocyte biology. 2018;104(3):447–59.

16. Girardin SE, Boneca IG, Viala J, Chamaillard M, Labigne A, Thomas G, et al. Nod2 is a general sensor of peptidoglycan through muramyl dipeptide (MDP) detection. The Journal of biological chemistry. 2003;278(11):8869–72.

17. Inohara N, Ogura Y, Fontalba A, Gutierrez O, Pons F, Crespo J, et al. Host recognition of bacterial muramyl dipeptide mediated through NOD2. Implications for Crohn's disease. The Journal of biological chemistry. 2003;278(8):5509–12.

18. Nachbur U, Stafford CA, Bankovacki A, Zhan Y, Lindqvist LM, Fiil BK, et al. A RIPK2 inhibitor delays NOD signalling events yet prevents inflammatory cytokine production. Nature communications. 2015;6:6442.

19. Tigno-Aranjuez JT, Benderitter P, Rombouts F, Deroose F, Bai X, Mattioli B, et al. In vivo inhibition of RIPK2 kinase alleviates inflammatory disease. The Journal of biological chemistry. 2014;289(43):29651–64.

20. Vieira SM, Cunha TM, França RF, Pinto LG, Talbot J, Turato WM, et al. Joint NOD2/RIPK2 signaling regulates IL-17 axis and contributes to the development of experimental arthritis. Journal of immunology (Baltimore, Md : 1950). 2012;188(10):5116–22.

21. GlaxoSmithKline. GSK2983559 First Time in Human Study. ClinicalTrials.gov, Identifier: NCT03358407.. 2017.

22. Wachtel MS, Shome G, Sutherland M, McGlone JJ. Derivation and validation of murine histologic alterations resembling asthma, with two proposed histologic grade parameters. BMC Immunol. 2009;10:58.

23. Haile PA, Votta BJ, Marquis RW, Bury MJ, Mehlmann JF, Singhaus R, Jr., et al. The Identification and Pharmacological Characterization of 6-(tert-Butylsulfonyl)-N-(5-fluoro-1H-indazol-3-yl)quinolin-4-amine (GSK583), a Highly Potent and Selective Inhibitor of RIP2 Kinase. J Med Chem. 2016;59(10):4867–80.

24. Zhang W, Lin C, Sampath V, Nadeau K. Impact of allergen immunotherapy in allergic asthma. Immunotherapy. 2018;10(7):579–93.

25. Shamji MH, Durham SR. Mechanisms of allergen immunotherapy for inhaled allergens and predictive biomarkers. J Allergy Clin Immunol. 2017;140(6):1485–98.

26. AstraZeneca. Study to evaluate Tezepelumab in Adults With Severe Uncontrolled Asthma. ClinicalTrials.gov, Identifier: NCT03927157.. 2019.

27. Liu YJ. Thymic stromal lymphopoietin: master switch for allergic inflammation. J Exp Med. 2006;203(2):269–73.

28. Froidure A, Shen C, Gras D, Van Snick J, Chanez P, Pilette C. Myeloid dendritic cells are primed in allergic asthma for thymic stromal lymphopoietin-mediated induction of Th2 and Th9 responses. Allergy. 2014;69(8):1068–76.

29. Chen ZG, Zhang TT, Li HT, Chen FH, Zou XL, Ji JZ, et al. Neutralization of TSLP inhibits airway remodeling in a murine model of allergic asthma induced by chronic exposure to house dust mite. PloS one. 2013;8(1):e51268.

30. Watanabe T, Minaga K, Kamata K, Sakurai T, Komeda Y, Nagai T, et al. RICK/RIP2 is a NOD2-independent nodal point of gut inflammation. Int Immunol. 2019;31(10):669–83.

31. Talreja J, Talwar H, Ahmad N, Rastogi R, Samavati L. Dual Inhibition of Rip2 and IRAK1/4 Regulates IL-1beta and IL-6 in Sarcoidosis Alveolar Macrophages and Peripheral Blood Mononuclear Cells. Journal of immunology (Baltimore, Md : 1950). 2016;197(4):1368–78.

32. Shaw PJ, Barr MJ, Lukens JR, McGargill MA, Chi H, Mak TW, et al. Signaling via the RIP2 adaptor protein in central nervous system-infiltrating dendritic cells promotes inflammation and autoimmunity. Immunity. 2011;34(1):75–84.

33. Jaafar R, Mnich K, Dolan S, Hillis J, Almanza A, Logue SE, et al. RIP2 enhances cell survival by activation of NF-kB in triple negative breast cancer cells. Biochem Biophys Res Commun. 2018;497(1):115–21.

34. Cai X, Yang Y, Xia W, Kong H, Wang M, Fu W, et al. RIP2 promotes glioma cell growth by regulating TRAF3 and activating the NFkappaB and p38 signaling pathways. Oncol Rep. 2018;39(6):2915–23.

35. Haffner CD, Charnley AK, Aquino CJ, Casillas L, Convery MA, Cox JA, et al. Discovery of Pyrazolocarboxamides as Potent and Selective Receptor Interacting Protein 2 (RIP2) Kinase Inhibitors. ACS Med Chem Lett. 2019;10(11):1518–23.

36. Haile PA, Casillas LN, Votta BJ, Wang GZ, Charnley AK, Dong X, et al. Discovery of a First-in-Class Receptor Interacting Protein 2 (RIP2) Kinase Specific Clinical Candidate, 2-((4-(Benzo[d]thiazol-5-ylamino)-6-(tert-butylsulfonyl)quinazolin-7-yl)oxy)ethyl Dihydrogen Phosphate, for the Treatment of Inflammatory Diseases. J Med Chem. 2019;62(14):6482–94.

37. Haile PA, Casillas LN, Bury MJ, Mehlmann JF, Singhaus R, Jr., Charnley AK, et al. Identification of Quinoline-Based RIP2 Kinase Inhibitors with an Improved Therapeutic Index to the hERG Ion Channel. ACS Med Chem Lett. 2018;9(10):1039–44.

38. Salla M, Aguayo-Ortiz R, Danmaliki GI, Zare A, Said A, Moore J, et al. Identification and Characterization of Novel Receptor-Interacting Serine/Threonine-Protein Kinase 2 Inhibitors Using Structural Similarity Analysis. J Pharmacol Exp Ther. 2018;365(2):354–67.

39. Goncharov T, Hedayati S, Mulvihill MM, Izrael-Tomasevic A, Zobel K, Jeet S, et al. Disruption of XIAP-RIP2 Association Blocks NOD2-Mediated Inflammatory Signaling. Mol Cell. 2018;69(4):551–65 e7.

40. Hrdinka M, Schlicher L, Dai B, Pinkas DM, Bufton JC, Picaud S, et al. Small molecule inhibitors reveal an indispensable scaffolding role of RIPK2 in NOD2 signaling. EMBO J. 2018;37(17).

41. Suebsuwong C, Pinkas DM, Ray SS, Bufton JC, Dai B, Bullock AN, et al. Activation loop targeting strategy for design of receptor-interacting protein kinase 2 (RIPK2) inhibitors. Bioorg Med Chem Lett. 2018;28(4):577–83.

42. Canning P, Ruan Q, Schwerd T, Hrdinka M, Maki JL, Saleh D, et al. Inflammatory Signaling by NOD-RIPK2 Is Inhibited by Clinically Relevant Type II Kinase Inhibitors. Chem Biol. 2015;22(9):1174–84.

43. Tigno-Aranjuez JT, Asara JM, Abbott DW. Inhibition of RIP2's tyrosine kinase activity limits NOD2-driven cytokine responses. Genes & development. 2010;24(23):2666–77.

