## Supplemental Figure 1 for "Immune Modulation of Allergic Asthma by Early Pharmacological Inhibition of RIP2"

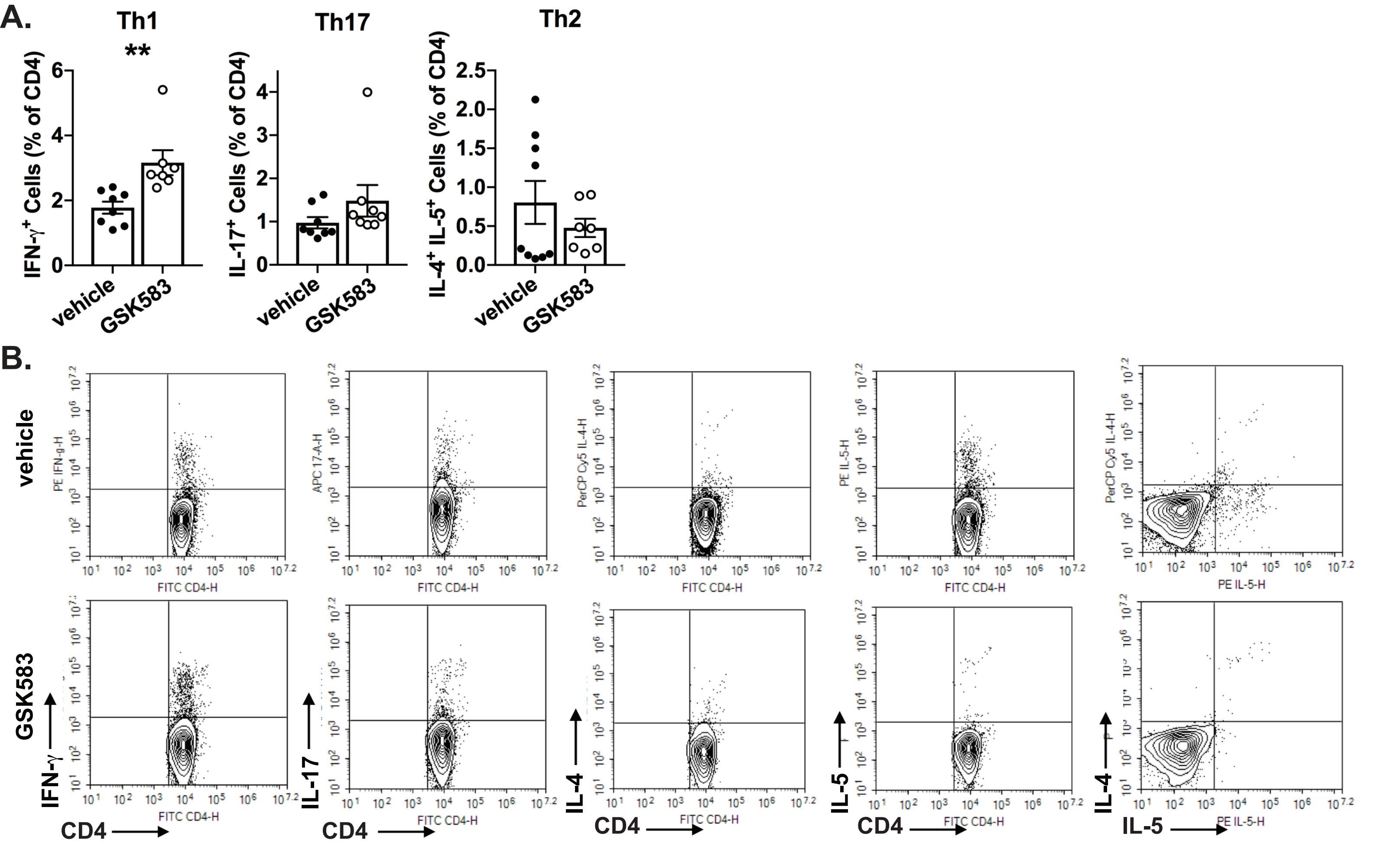


**Supplemental Figure 1.** Pharmacologic inhibition of RIP2 downregulates Th2 immunity and upregulates Th1 immunity in a HDM model of asthma. WT C57BL/6 mice were subjected to an acute HDM asthma model and were administered either regular chow or chow containing RIP2 inhibitor (GSK583) as indicated in Fig1A. On day 14, mice were euthanized and lung draining lymph node cells were stimulated with PMA and ionomycin and subjected to intracellular cytokine staining (ICS). A.) The percentage of $CD4^{+}IFN\text{-}\gamma^{+}$(Th1) , $CD4^{+}IL\text{-}17^{+}$(Th17), or $CD4^{+}IL\text{-}4^{+}IL\text{-}5^{+}$(Th2) are shown for vehicle compared to GSK583-treated mice. B.) Flow cytometry gating strategy for ICS percentages shown in A.). C.) Levels of HDM-specific total IgG and $IgG_1$ antibodies in the serum of vehicle chow or RIP2 inhibitor (GSK583) chow-treated mice subjected to an acute HDM asthma model as measured by ELISA. Data are presented as scatterplots with bars where bar heights represent means ± SEM. Filled in circles represent individual datapoints for vehicle/control chow-treated mice and open circles represent individual datapoints for GSK583-treated mice. Data are pooled from 3 independent experiments for a total of n=9 mice per group. Statistical analysis was performed using an unpaired, two-tailed Student's t-test. **= $p<0.01$
